# Brain capillary obstruction as a novel mechanism of anti-CD19 CAR T cell neurotoxicity

**DOI:** 10.1101/2021.05.25.445614

**Authors:** Lila D Faulhaber, Kendra Jae Hartsuyker, Anthea Q Phuong, Yeheun Cho, Katie K Mand, Stuart D Harper, Aaron K Olson, Gwenn A Garden, Andy Y Shih, Juliane Gust

## Abstract

Immunotherapy for hematologic malignancies with CD19-directed CAR T cells is associated with neurotoxicity in about 40% of patients. Systemic cytokine release syndrome, endothelial activation, and disruption of endothelial integrity have all been associated with neurotoxicity, but it remains unclear how these mechanisms interact and how they lead to neurologic dysfunction. We developed a syngeneic mouse model which manifests systemic cytokine release and behavioral abnormalities within 3-5 days after infusion of high-dose murine CD19-CAR T cells. Histologic examination revealed widespread brain hemorrhages, diffuse extravascular IgG deposition, loss of capillary pericyte coverage and increased prevalence of string capillaries. *In vivo* two-photon imaging of blood flow revealed plugging of >10% of capillaries by leukocytes, associated with regions of localized hypoxia. These data reveal capillary obstruction and associated brain hypoxia and microvascular decline as a potential basis for neurotoxicity during CD19-CAR T cell treatment in humans, which may be amenable to therapeutic interventions.

## Introduction

Chimeric antigen receptor (CAR) T cell therapy has been revolutionizing the treatment of hematologic malignancies, but systemic and neurologic toxicities remain a major concern^1^. CAR T cells work by recognizing surface targets on tumor cells via a chimeric receptor that consists of an extracellular binder, typically an antibody fragment, fused to intracellular signaling domains^2^. Once the CAR T cells are infused into the patient, they recognize and lyse the target cells via formation of a nonclassical immune synapse^3^. Binding of targets also induces CAR T cell proliferation and systemic cytokine release. The greatest success of this therapeutic approach is seen in CD19^+^ hematologic malignancies such as acute lymphocytic leukemia and non-Hodgkin’s lymphoma, with remission rates above 90% and FDA approval of multiple CD19-targeting CAR T products^4^.

Despite these successes, neurologic toxicity remains a major concern in CAR T cell therapy. Neurotoxicity, also termed Immune Effector Cell Associated Neurotoxicity (ICANS)^5^ occurs in approximately 40% of patients receiving CD19-directed CAR T cells, and is also seen after CD22- and B cell maturation antigen (BCMA)-CAR treatment^6,7^. The most common symptoms are transient language disturbance or delirium, typically accompanied by EEG background slowing^8–11^. Approximately 10-20% of patients have potentially life-threatening manifestations such as seizures and coma. Empiric treatments include corticosteroids and IL-1 blockade, which is currently being evaluated in multiple clinical trials^12^. Unfortunately, ~1% of patients treated with CD19-directed CAR T cells develop rapidly progressive cerebral edema that can lead to death despite aggressive immunomodulation and intracranial pressure-directed interventions^6^. Autopsy findings in patients with cerebral edema include microhemorrhages, microvascular disruption and perivascular plasma extravasation, perivascular infiltration of T cells and macrophages, leptomeningeal accumulation of CAR T cells, and astrocyte activation^7,13^.

The mechanism of neurotoxicity remains incompletely understood. The most consistent clinical risk factor is the severity of cytokine release syndrome (CRS), which in turn is influenced by disease burden and amount of CAR T cell proliferation^12,14–16^. Dysfunction of the neurovascular unit likely plays a role, as evidenced by reversible interstitial edema on MRI in some patients^8–10^, and a shift of the angiopoietin-Tie2 axis toward endothelial activation that was demonstrated in several clinical studies^8–10^. The most faithful animal model to date has been in nonhuman primates, which develop CRS and neurotoxicity after treatment with CD20-directed CAR T cells^17^. In mice, no model has yet been described that recapitulates multiple key features of neurotoxicity, including time course, behavioral changes, and/or histopathologic findings similar to those in humans^7^. In immunodeficient mice treated with human CAR T cells, some aspects of neurotoxicity have been modeled and linked to systemic cytokine release. In a humanized mouse model of CD19-CAR T toxicity, blockade of IL-1 prevented neurologic deterioration 4 weeks after CAR T cell infusion^18^. In a xenograft mouse model, blockade of granulocyte-macrophage colony-stimulating factor (GM-CSF) reduced contrast extravasation on brain MRI^19^. Most syngeneic mouse models, where murine CAR T cells are given to immunocompetent mice, have shown excellent CAR T cell function but no toxicity^20–24^. In the syngeneic studies that did demonstrate toxicity, detailed neurologic phenotyping has not yet been reported^25–27^. We have developed an immunocompetent mouse model that develops CRS and neurotoxicity after high dose cyclophosphamide lymphodepletion and high dose second-generation murine CD19-directed CAR T cells. We show evidence of blood-brain-barrier disruption, pericyte injury and capillary regression. By *in vivo* two-photon imaging of cerebral cortex, we found persistent capillary plugging by leukocytes and tissue hypoxia, which supports a key role for microvascular dysfunction in the pathophysiology of neurotoxicity.

## Results

### CD19-CAR T cell treatment induces cytokine release syndrome and neurotoxicity

To explore the mechanism of neurotoxicity during CAR T cell treatment, we used a syngeneic mouse model of CD19-CAR T cell treatment (Fig. 1a,b). To avoid confounding effects from tumor growth or lysis, we conducted all experiments in nontumor bearing wild type BALB/c mice, where CD19-CAR T cells target CD19 on normal B cells^28^. Control mice received syngeneic T cells which underwent mock transduction without viral vector (“mock T cells”). All mice received cyclophosphamide for lymphodepletion one day prior to treatment. CD19-CAR T cells expanded well despite absence of a tumor target (Fig. 1c) and remained detectable for at least 56 days (data not shown). CAR T cells eradicated B cells as expected (Fig. 1d), and B cell aplasia also persisted for at least 56 days (data not shown). Concurrently with the rapid expansion of the CAR T cells, mice developed dose-dependent hypothermia (Fig. 1e) and weight loss (Fig. 1f). We did not observe any fevers, but some animals with severe toxicity developed hypothermia below 34 degrees C.

**Figure 1.**
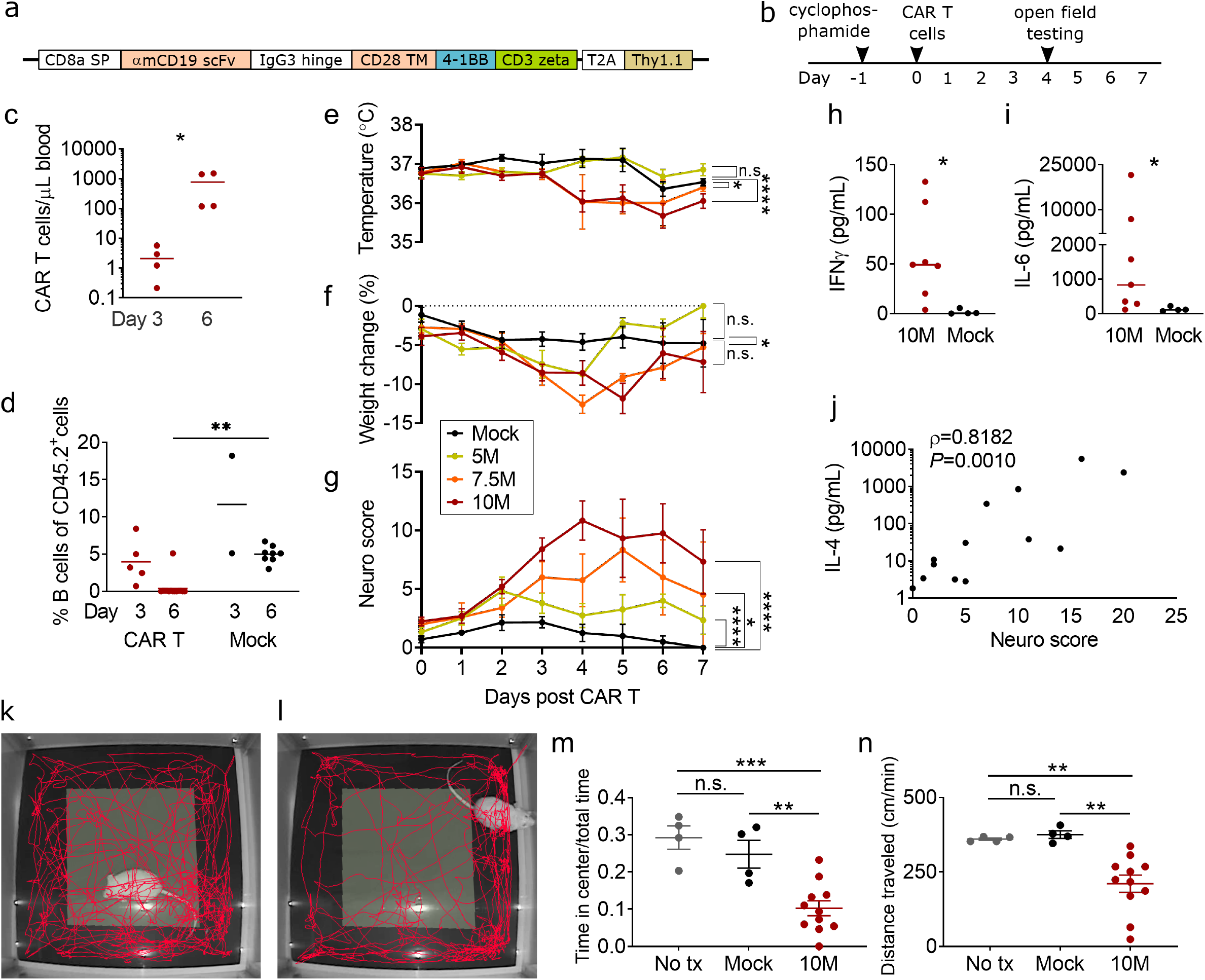
Neurotoxicity and CRS after CAR T cell treatment. **a**, structure of the murine CD19-directed CAR. SP, signal peptide; TM, transmembrane domain. **b**, experimental scheme. **c**, CAR T cells (CD3^+^ Thy1.1^+^) in blood by flow cytometry. X-axis denotes day after CAR T cell injection, 5-10 x 10^6^ cells/mouse. **d**, CD19^+^ B cells as fraction of total CD45^+^ splenocytes. Depletion by CAR T cells is complete by day 6. **e**, ear temperature, **f**, body weight, and **g,** neurologic exam scores in the first 7 days after CAR T cell infusion, P values were calculated by linear mixed models (mixed-effects analysis). **h**, serum levels of interferon-gamma on day 4 after treatment with 10 million CAR T cells (10M) or control T cells (Mock). **i**, serum levels of IL-6. **j**, correlation of IL-4 serum levels and same-day neuro score, Spearman’s r. **k,l**, representative examples of 5-minute tracks in the open field for a CAR T cell treated (k) and mock T cell treated (l) mouse. **m,n**, open field test, y-axis denotes the fraction of time spent in the center of the arena (m) and locomotion speed (l) during the 5 minute test. c,d,m,n: lines show the mean, statistics by one-way ANOVA with Holm-Sidak posttest. h, i: lines show the median, Mann-Whitney test. All panels: * P<0.05, ** P<0.01, *** P<0.001, **** P<0.0001, all other comparisons are P>=0.05. Each data point denotes one animal, all data pooled from 3 or more independent experiments.

To measure behavioral changes consistent with neurotoxicity, we adapted a 20-item test (“neuro score”) based on a neurophenotyping scoring system that was developed for mutant mouse screening^29^. Doses of 5 x 10^6^ CAR T cells induced only mild elevations in neuro score, whereas 7.5 x 10^6^ and especially 10 x 10^6^ CAR T cells per mouse induced severe behavioral abnormalities that peaked on day 4-5 after CAR T cell administration (10 x 10^6^ CAR T cells vs mock, *P*<0.0001, mixed-effects analysis) (Fig. 1g). There was no effect of sex or age on peak neuro score within any of the dose groups (data not shown). The most frequent abnormalities on the neuro score were deficits in postural adjustment, balance, and exploration. In addition, mice frequently had piloerection, hunched posture, and alterations in breathing patterns (fast breathing in milder toxicity, and deeper, more labored breaths during severe toxicity). We did not observe seizures or focal motor dysfunction.

Since the clearest phenotype was present in the mice receiving 10 million CAR T cells, we focused on this dose level in subsequent experiments. The behavioral abnormalities were accompanied by elevations in serum cytokines. On day 4-5 after CAR T cell infusion, serum levels of IFNγ, IL-2, IL-4, IL-5, IL-6, and IL-10 were significantly elevated in mice receiving 10 x 10^6^ CAR T cells, compared to mice who received mock transduced CAR T cells (Fig. 1 h,i; Table 1). We observed marked heterogeneity of serum cytokine levels between individual CAR T treated mice, even though all animals had detectable CAR T cells and B cell suppression. Since neurotoxicity scores also varied between individuals, we hypothesized that differences in systemic proinflammatory cytokine profiles could account for the differences in observed neurological phenotype. Indeed, linear regression analysis showed a positive association of neurologic exam scores and serum levels of IL-4 (Fig. 1j), IL-5, and IL-10, while there was no correlation of neuro score with serum levels of CXCL1, IL1β, IL-2, IL-6, IL-12p70/IL-25, IFNγ, or TNF (Table 1). Open field testing on day 4 after CAR T cell infusion showed that mice treated with 10 x 10^6^ CAR T cells had markedly decreased locomotion speed (*P*=0.0060 vs mock) and spent a smaller fraction of time in the center of the field (*P*=0.0043 vs mock) (one-way ANOVA with Holm-Sidak posttest to correct for multiple comparisons) (Fig. 1 k-n). This indicates that CAR T treated mice did not only show a decrease in activity level, but also an increase in avoidant or anxious behavior.

**Table 1.** Serum cytokine levels in non-tumor bearing mice after CAR T cell treatment. Left column shows day 4+5 measurements from mice receiving 10 million CAR T cells compared to mock controls. Right column shows the correlation of day 4-7 serum cytokine levels with same-day neuro scores in mice receiving 5-10 million CAR T cells.

|  | **10M CAR T**<br>**(N=7)** | **Mock**<br>**(N=4)** | *P* (Mann-Whitney) | **Correlation with severity of neurotoxicity (5-10M CAR T)**<br>**(N=14)** |  |
| --- | --- | --- | --- | --- | --- |
|  | Median<br>pg/mL | Median<br>pg/mL |  | Pearson's R | *P* (two-tailed *t*<br>unpaired test) |
| **CXCL1** | 731.6 | 386.4 | 0.2309 | **0.6425** | **0.0179** |
| **IFNγ** | **49.3** | **0.6** | **0.0121** | 0.0249 | 0.9356 |
| **IL-1β** | 18.8 | 2.396 | 0.2303 | 0.1313 | 0.6689 |
| **IL-2** | **3.0** | **0.8** | **0.0242** | 0.0005382 | 0.9986 |
| **IL-4** | **342.8** | **0** | **0.0061** | **0.6585** | **0.0144** |
| **IL-5** | **2723** | **13.6** | **0.0242** | **0.6822** | **0.0102** |
| **IL-6** | **836.4** | **112.2** | **0.0121** | **0.5715** | **0.0413** |
| **IL-10** | **239.4** | **5.1** | **0.0121** | **0.7659** | **0.0023** |
| **IL-12p70** | 0 | 0 | 0.6606 | **0.5668** | **0.0434** |
| **TNF** | 132.7 | 43.5 | 0.1091 | 0.06023 | 0.845 |

Taken together, these data show that wild-type non tumor bearing mice develop acute cytokine release syndrome and abnormal behavior after treatment with CD19-directed CAR T cells. The timing of the behavioral changes coincides with the period of CAR T cell proliferation and cytokine release, similar to what occurs in human patients.

### Mice with neurotoxicity develop cerebral microhemorrhages and focal endothelial disruption

In humans, severe CAR T neurotoxicity can be accompanied by endothelial activation and injury to the cerebral microvasculature.^6^ We examined the brain tissue in our mouse model to determine if similar microvascular changes were occurring. Indeed, we noted cerebral microhemorrhages in the mice receiving 10 x 10^6^ CAR T cells per mouse) (mean 64, range 17-140 hemorrhages per brain, *P*=0.0188, 2-tailed one sample t test) (Fig. 2a,b). Hemorrhages occurred in all brain regions, including cortex, basal ganglia, hypothalamus, brainstem, and cerebellum (Fig. 2). No hemorrhages were seen in mice treated with mock transduced T cells, or those receiving the lower doses of 5×10^6^ or 7.5×10^6^ CAR T cells per mouse.

**Figure 2.**
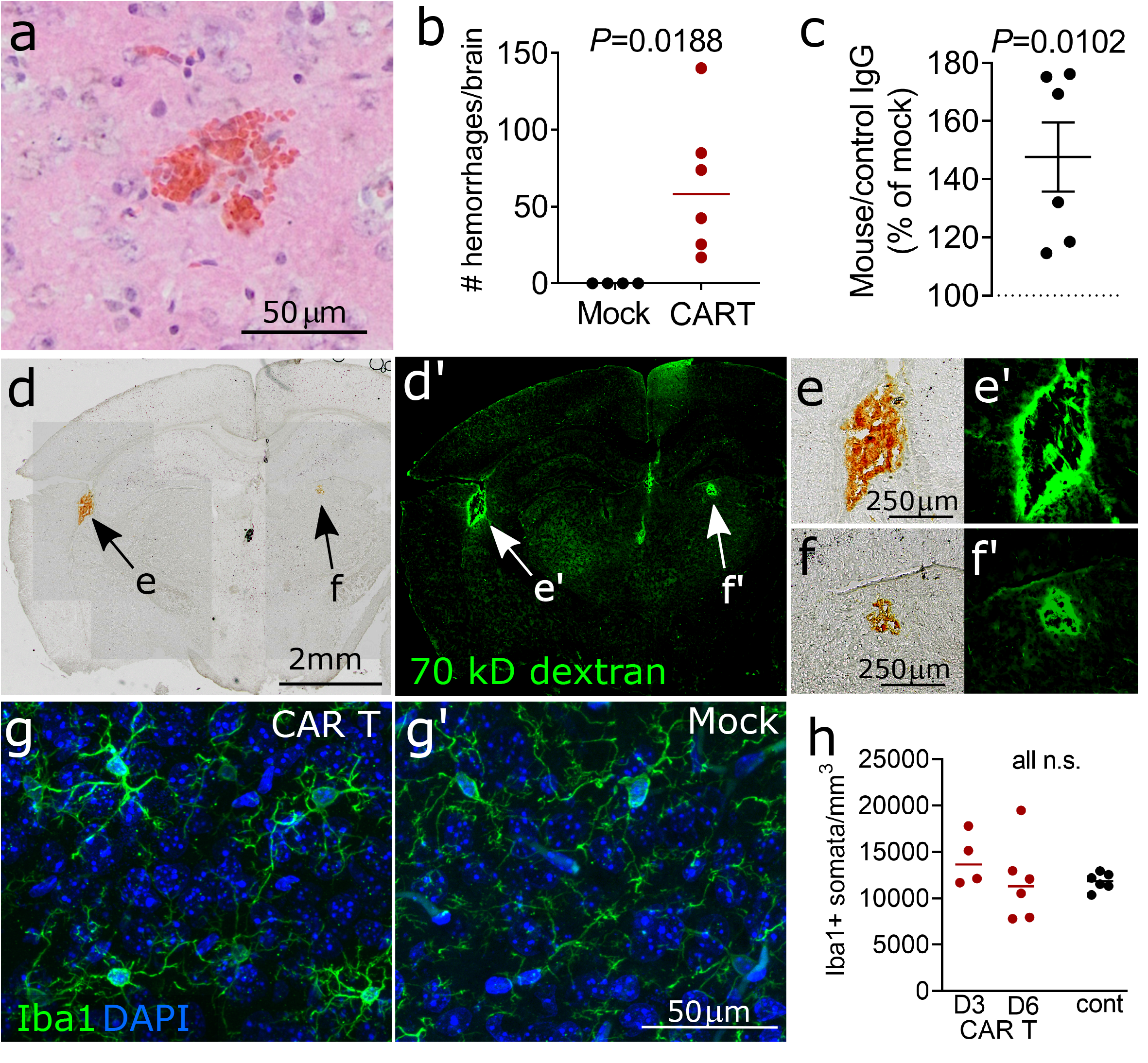
Cerebral microvascular injury during neurotoxicity. **a**, brightfield image of H&E stained brain section showing acute microhemorrhage in a mouse that received 10 x 10^6^ CAR T cells 4 days prior. **b**, quantification of brain microhemorrhages in the entire brain. **c**, quantification of anti-mouse IgG immunofluorescence normalized to control species IgG immunofluorescence. Each data point shows median of 3 brain sections from one mouse, normalized to concurrently treated controls receiving mock transduced T cells. CAR T cell treated mice received 5, 7.5, or 10 x 10^6^ CAR T cells 4-6 days prior to sacrifice. **d**, bright field image of coronal brain section from a CAR T treated mouse that received i.v. FITC-dextran 30 minutes prior to sacrifice. Arrows indicate microhemorrhages, which are visible without additional tissue staining. **d’** shows fluorescence imaging of the same section, demonstrating FITC-dextran extravasation surrounding the hemorrhages. **e,** magnification of subcortical hemorrhage in c, **f**, magnification of basal ganglia hemorrhage. **e’** and **f’** show the corresponding fluorescence images. **g, g’**: representative immunofluorescence of cortical microglia labeled with anti-Iba1 antibody. In the CAR T treated mouse (**g**), the microglia maintain a similar finely ramified resting appearance as mock controls (**g’**). All graphs: each data point represents one mouse, pooled data from 3 experiments, lines show mean. **b, c**: one-sample 2-tailed t test. **h**, one-way ANOVA with Holm-Sidak posttest.

To determine whether more widespread blood-brain-barrier disruption was present even in areas without microhemorrhages, we used immunohistochemistry to quantify IgG deposition in the cortex as a measure of increased permeability to macromolecules^30^. CAR T cell treated mice had a mean of 47% higher levels of immunofluorescence against mouse IgG compared to mock treated controls (P=0.0102, one-sample t test) (Fig. 2c). As another measure of blood-brain barrier disruption, we injected 70kDa fluorescent-conjugated dextran 30 minutes prior to sacrificing the mice. We observed extensive leakage of dextran around microhemorrhages, but there was no visible extravasation of dextran in areas without hemorrhage (Fig. 2d-f). Some amount of fluorescently labeled dextran remained visible within the cerebral microvessels even after thorough transcardiac perfusion. This precluded the use of quantification of overall brain dextran fluorescence for measuring blood-brain-barrier permeability, since we would not have been able to distinguish between intravascular and extravascular dextran.

Since activation and/or proliferation of brain resident microglia occurs during many neuroinflammatory conditions^31^, we reasoned that microglial activation may accompany the proinflammatory cytokine environment. Surprisingly, we found normal microglial ramification on histologic examination, as is indicative of a quiescent surveillance state (Fig. 2g). There was no difference in the number of Iba^+^ microglia either on day 3-4 (median 13,649 cells/mm^3^) or day 6-7 after CAR T cells (median 11,310 cells/mm^3^) compared to mock treated control (11,861 cells/mm^3^, *P*=0.4269) (Fig. 2h).

### Loss of pericyte coverage and increased incidence of string capillaries

Given the observation of microhemorrhages and increased IgG deposition in the brain, we hypothesized that components of the blood-brain barrier might show morphologic abnormalities even in areas without frank hemorrhage. Claudin-5, a key component of brain endothelial tight junctions^32^, had unchanged appearance on immunofluorescence on day 6 after CAR T cell treatment (Fig. 3a). The capillary endothelial basement membrane also appeared intact, as evidenced by laminin^33^ immunofluorescence which did not show gaps or breaks (Fig. 3b), and covered the same capillary lengths per volume of cortex in CAR T (573 mm/mm^3^) and mock treated (581 mm/mm^3^) groups (*P*=0.9329, unpaired t test). However, pericyte coverage of cortical capillaries trended toward decrease in CAR T treated animals (Fig. 3c,d). Although the mean uncovered capillary length was not statistically significantly different (8.7% in animals receiving 10 x 10^6^ CAR T cells, compared to 3.9% in mock controls, *P*=0.1200), the phenotype was heterogeneous among animals (Fig. 3e). To better understand why some CAR T treated mice had more uncovered capillaries than others, we compared the neurologic symptom score to pericyte coverage, and found that mice with more severe neurologic phenotypes also had a greater fraction of uncovered capillaries (Spearman ρ = 0.6216, *P*=0.0262, Fig. 3f). This did not appear to be due to loss of the pericytes themselves, since the numbers of pericyte cell bodies were similar both as measured by capillary length (CAR T=14.1 and mock=9.9 pericytes per millimeter capillary length, *P*=0.1095, unpaired t test) or by volume of brain tissue (CAR T mean=7839 and mock mean=5766 pericytes per mm^3^, *P*=0.1819, unpaired t test). Since pericyte coverage of microvessels is important for maintaining capillary network structure^34^, we also searched for evidence of capillary regression. Histologically, involuting microvessel segments can have the appearance of string capillaries, which consist of basement membrane but lack a patent lumen^35^. Indeed, string capillaries were significantly more common in CAR T treated animals (mean 249 string capillaries/mm^3^) compared to mock (46 string capillaries/mm^3^) (*P*=0.0280, unpaired t test) (Fig. 3g,h).

**Figure 3.**
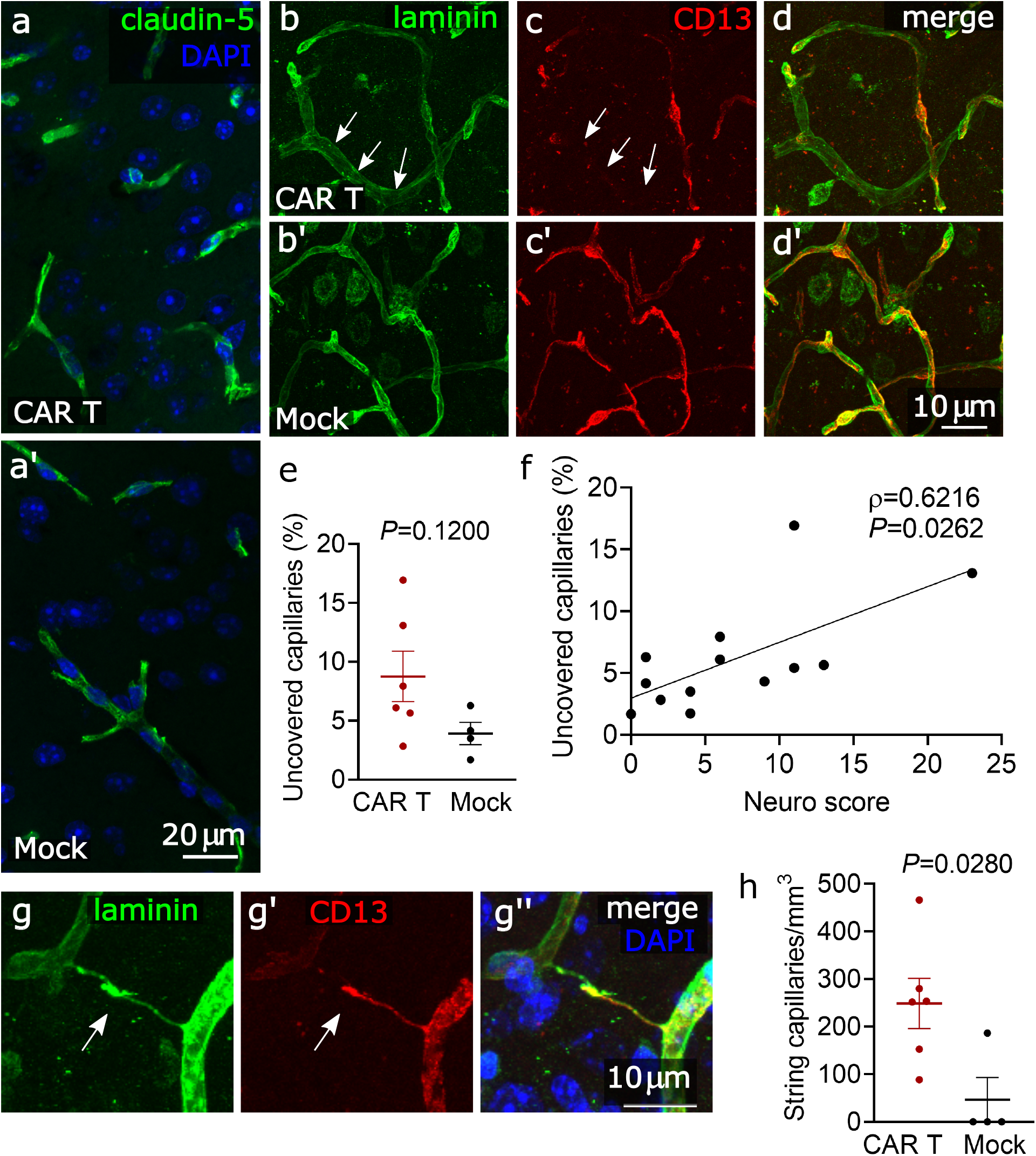
Pericyte disruption and string capillaries. **a**, representative images of claudin-5 immunofluorescence showing no deficit in endothelial tight junctions in a mouse treated with 10 x 10^6^ CAR T cells 6 days prior, compared to mock T cell treated control. **b**, representative images of laminin immunofluorescence showing intact capillary endothelial basement membrane in a mouse treated with 10 x 10^6^ CAR T cells 6 days prior (b) compared to mock T cell treated control CAR T (b’). **c**, CD13 immunolabeling showing pericytes in same areas as b/b’. Arrows indicate a capillary with loss of pericyte coverage. **d**, overlay of laminin and CD13 immunolabeling in same areas as b/b’. **e**, capillary pericyte coverage in mice receiving 10 x 10^6^ CAR T cells compared to mock treated mice, unpaired t test. **f**, correlation of pericyte coverage and neurological exam scores for mice receiving 0-10 x 10^6^ CAR T cells. r, Spearman rank correlation. **g**, representative image of a string capillary lacking a lumen (arrow). **h**, string capillary counts per mm^3^ of cortex in mice receiving 10 x 10^6^ CAR T cells compared to mock treated mice, unpaired t test. All images are from somatosensory cortex. All graphs show pooled data from 3 experiments, each data point represents one mouse, error bars show mean +/- SEM.

### Brain capillary plugging during neurotoxicity

We next sought to determine whether the abnormalities seen on histology correspond to functional changes in the perfusion of the cerebral microvasculature. To measure cerebral capillary blood flow, we performed *in vivo* two-photon imaging during the period of the most severe behavioral neurotoxicity (days 4 and 6) (Fig. 4a). We used a thinned-skull cranial window preparation to minimize surgically induced inflammation of the underlying meninges and brain (Fig. 4b). We acquired ~150μm deep z-stacks of somatosensory cortex to analyze the anatomy and flow patterns of the microvasculature. Each z-stack was imaged prior to CAR T cell treatment and on day 4 and 6 post treatment. Strikingly, 5.4% of cortical capillaries were nonflowing on day 4, which appeared to be due to cells plugging the capillaries (Fig. 4c-f). We only considered plugs lasting more than 5 seconds, since momentary stalls can be caused by large leukocytes even in normal conditions. On day 6, the fraction of plugged capillaries increased to 11.9%, compared to 1.1% in controls (*P*=0.0072) (Fig. 4g). This level of capillary plugging is very severe and likely to cause behavioral abnormalities, as only 2% capillary plugging was sufficient to cause cognitive abnormalities in a mouse model of Alzheimer’s Disease^36^. The abnormal capillary blood flow did not appear to be related to poor systemic perfusion or oxygenation: mean peripheral oxygen saturation measured by paw infrared sensor during imaging was 86% (range 75-91%) vs. 85% in mock controls (range 79-95%), and mean heart rate was 479 beats per minute (range 416-545) vs. 475 in mock controls (range 312-542). Echocardiograms showed preserved cardiac output (mean 13.3ml/min in CAR T, 14.5ml/min in mock on day 4, *P*=0.7506; 18.2ml/min CAR T, 16.7ml/min mock on day 6, *P*=0.8583; unpaired t test; N=5 CAR T, N=2 mock).

**Figure 4.**
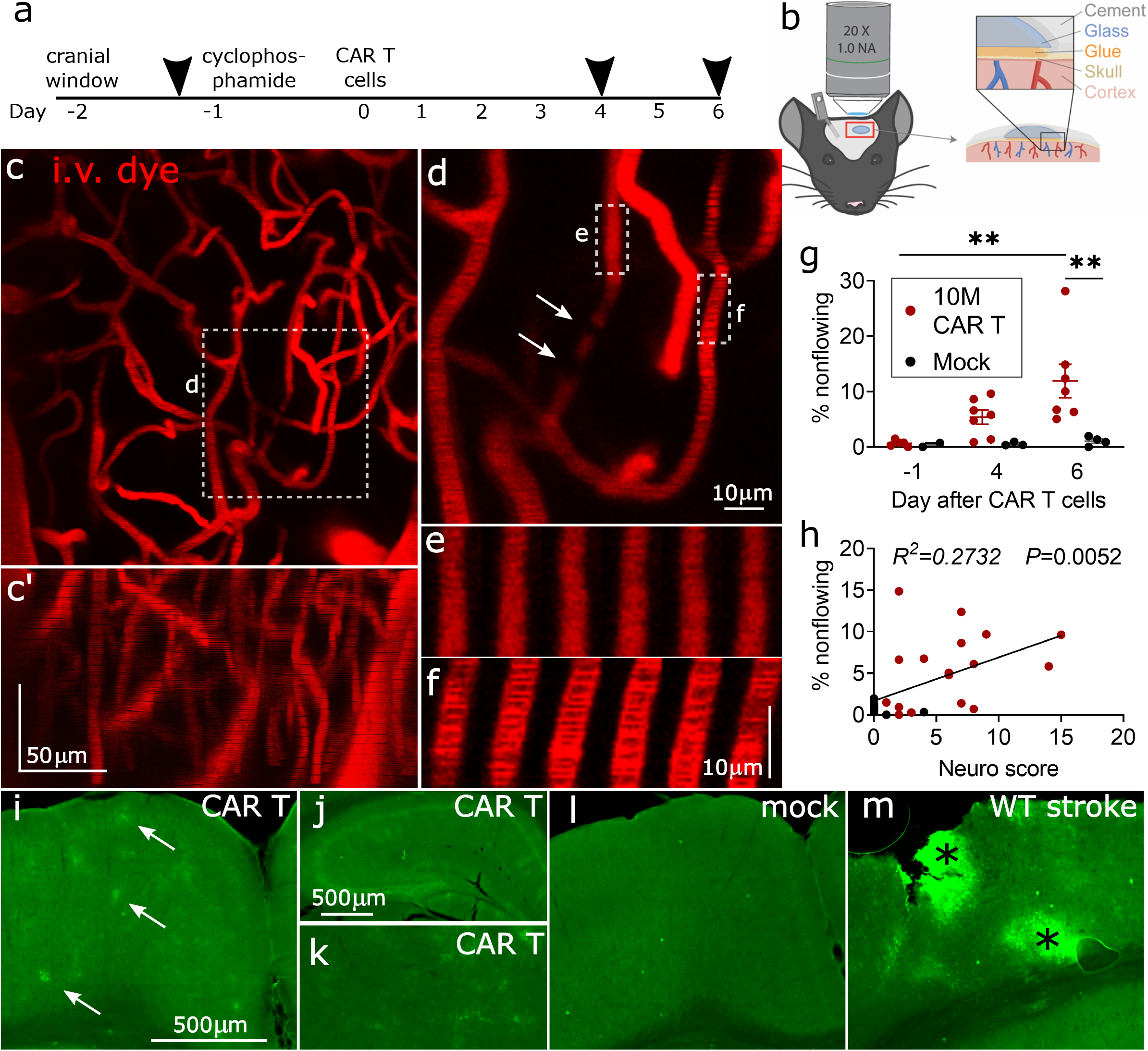
Cortical capillary plugging visualized by *in vivo* two photon imaging. **a**, timeline of experiments, arrowheads denote imaging sessions. **b**, schematic of thinned skull *in vivo* imaging preparation. timeline of experiments **c**, representative image of cortical capillary bed, z-projection viewed from top down, and **c’** from the side, with the top of the image oriented toward the pial surface of the cortex. **d**, close-up of a nonflowing capillary with plugs (arrows) that are seen as filling defects. A section of a nonflowing (**e**) and a flowing (**f**) capillary is shown as a montage of 6 successive scans. The images are separated by 1.14 seconds in time and 1mm in the z axis. In the flowing capillary in **f**, moving shadows of red blood cells can be seen, which are absent in **e**. **g**, percentage of nonflowing cortical capillaries during each of the imaging time points. **P<0.01, one-way ANOVA with Holm-Sidak posttest, bars show mean +-SEM. Each data point represents one animal, data from three separate experiments. **h**, correlation of behavioral neuro score on day of imaging and percentage of nonflowing capillaries, Pearson’s R. **i-m**, hypoxyprobe immunofluorescence in coronal brain sections, pial surface is at the top: **i**, patches of hypoxic cortex in a CAR T cell treated mouse (arrows). Patchy hypoxia is also visible in hippocampus (**j**, note different scale bar), and thalamus (**k**). In mock treated control cortex, there is no such patchy hypoxyprobe labeling (**l**). **m**, positive control – a small stroke was made by laser coagulation of a cortical venule, resulting in two areas of hypoxia (asterisks).

Consistent with capillary plugging being a contributor to behavioral abnormalities, neuro scores were higher in mice with more severe capillary plugging (*P*=0.0052, Pearson’s correlation)(Fig. 4h). Further supporting the pathologic effect of capillary plugging, hypoxyprobe immunolabeling was abnormal in CAR T cell treated mice. Patches of hypoxic cells were apparent throughout the brain, including cortex, but also in deeper regions such as thalamus and hippocampus (Fig. 4i-k). No hypoxia labeling was seen in control mice (Fig. 4l). Specificity of hypoxyprobe labeling was confirmed in a positive stroke control (Fig. 4m).

### Capillary plugs are leukocytes

To identify the cause of the capillary plugging, we first used i.v. injections of Rhodamine 6G^37^, which labels leukocytes and platelets. Capillary plugs were consistently Rhodamine 6G positive (Fig. 5a). To further confirm the identity of the plugs, we injected fluorescently conjugated antibodies during imaging (anti-CD45.2 as a pan-leukocyte label, and anti-CD3 to identify T cells). In CAR T treated mice, but not in mock controls, we observed large numbers of crawling or firmly adherent CD45.2+ leukocytes in pial venules, a subset of which was also positive for CD3 (Fig. 5b). All capillary plugs labeled with CD45.2, and many were double positive for CD45.2 and CD3 (Fig. 5c,d). Given the observation of increased crawling, adhesion, and capillary stalling by T cells and other leukocytes, we next asked whether CAR T cell treatment leads to increased migration of peripheral leukocytes into the brain parenchyma. Flow cytometry of enriched brain leukocytes showed no significant increase of immune cell migration in CAR T cell treated mice (Fig. 5e-g) (median 158 CD45^hi^ cells/mg brain tissue) compared to mock control (median 76 CD45^hi^ cells/mg) (*P*=0.5697, Mann-Whitney test). The number of infiltrating T cells was also not elevated, with 3.6 CD3+ cells per mg brain tissue compared to 1.5 CD3+ cells in controls (*P*=0.7758, Mann-Whitney test) (Fig. 5h). CAR T treated mice had a median of just 1.4 CAR T cells/mg brain (Fig. 5i). Taken together, these data support the theory that intravascular obstruction by leukocytes is a key event in the pathogenesis of CAR T neurotoxicity.

**Figure 5.**
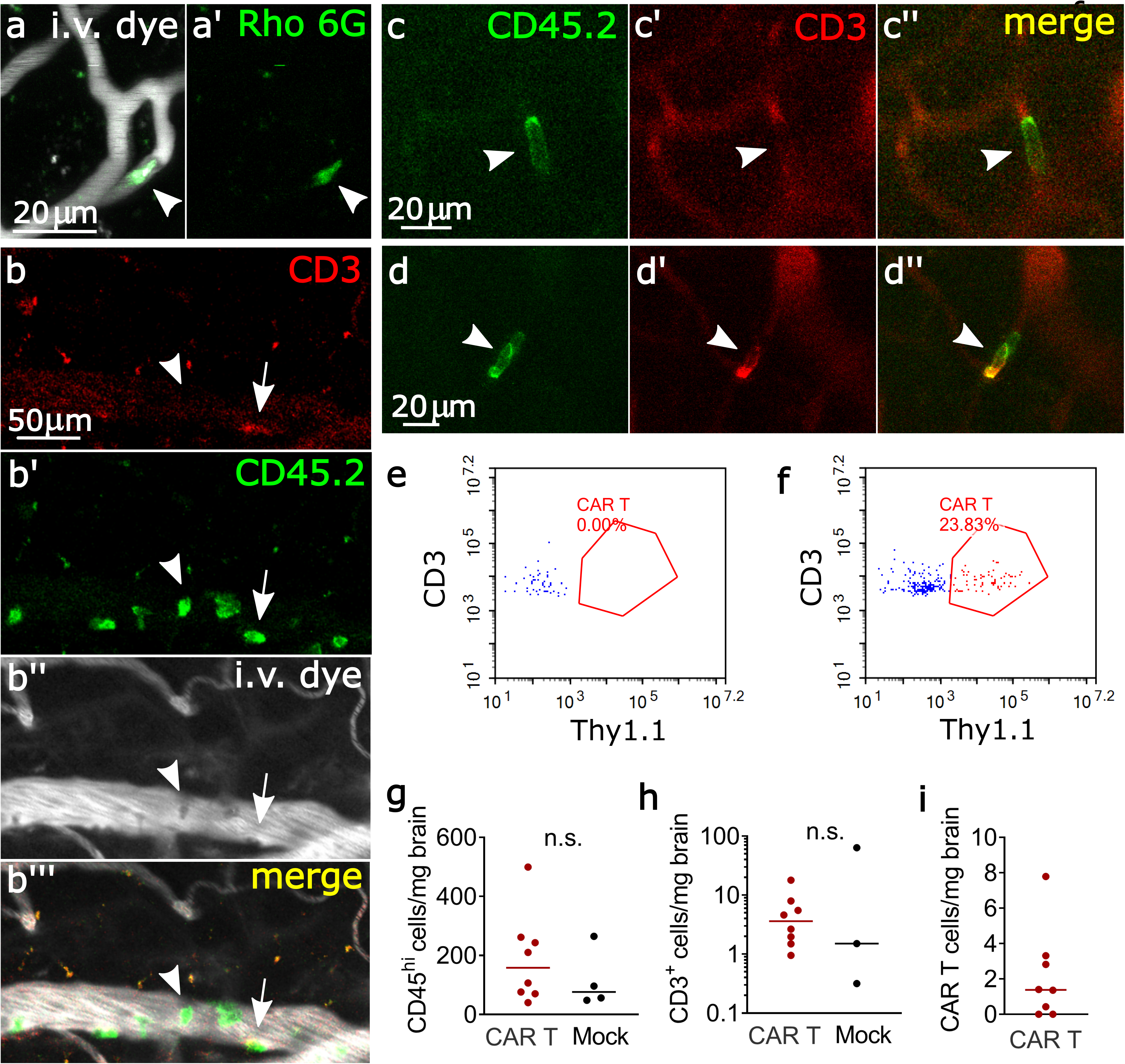
Leukocyte plugging of cortical capillaries. **a**, representative image of a Rhodamine 6G positive capillary plug. **b**, pial venule with multiple leukocytes adhering to the endothelium. Most were CD45.2^+^, CD3^-^ (arrowhead), with scattered T cells (CD45.2^+^, CD3^+^, arrow) **c**, CD45.2^+^, CD3^-^ leukocyte (arrowhead) that remained stationary in a capillary over a 3-minute time scan. **d**, CD45.2^+^, CD3^+^ T cell plugging a capillary. In **c** and **d**, vessels are outlined in the red channel by circulating unbound anti CD3-Alexa 594 antibody.). **e,f**, representative flow cytometry plots of brain infiltrating untransduced T cells (blue) and CAR T cells (red) in mice treated with mock (**e**) and CAR (**f**) T cells. Thy1.1 is the transduction marker coexpressed with the CAR. **g-i**, flow cytometry of brain-infiltrating leukocytes on day 3-6 after treatment. Each data point represents one mouse, mice received 5-10 x 10^6^ CD19 CAR T cells or 10 x 10^6^ mock transduced T cells, pooled data from 3 replicate experiments. Lines show the median. Mann-Whitney test.

### Capillary diameter and network structure during neurotoxicity

We next considered the role of capillary diameter in the etiology of capillary obstruction by leukocytes. To determine if simple mechanical obstruction by enlarged leukocytes could cause plugging, we measured the baseline (pre-treatment) diameters of the capillaries that later developed plugs. Only capillaries with a diameter of less than 5.5μm developed plugs (Fig. 6a). This is consistent with our observation that all plugging objects are leukocytes, which are likely too small to obstruct any of the larger vessels. If simple mechanical obstruction was the cause of plugging, then the smaller a capillary, the more likely it should end up being plugged by a circulating leukocyte. But the likelihood of plugging remained similar in the size range from 2.5 to 4.5μm (Fig. 6a), and leukocytes were able to deform to fit through small capillaries (Fig. 5c,d). We also considered the possibility that systemic cytokine release could globally decrease capillary diameters, thereby increasing the fraction of capillaries smaller than 5.5μm and increasing the likelihood of plugging. However, there was no statistically significant difference in capillary diameters on days −1, 4 and 6 (Fig. 6b,c), diameter change between imaging days (Fig. 6d,e), or variance of diameter changes (data not shown) between CAR T and mock treated mice. These data support the conclusion that increased leukocyte adhesion to the endothelium is required for capillary plugging to occur during CAR T neurotoxicity, and that the phenotype could be reversed by decreasing leukocyte adhesion.

**Figure 6.**
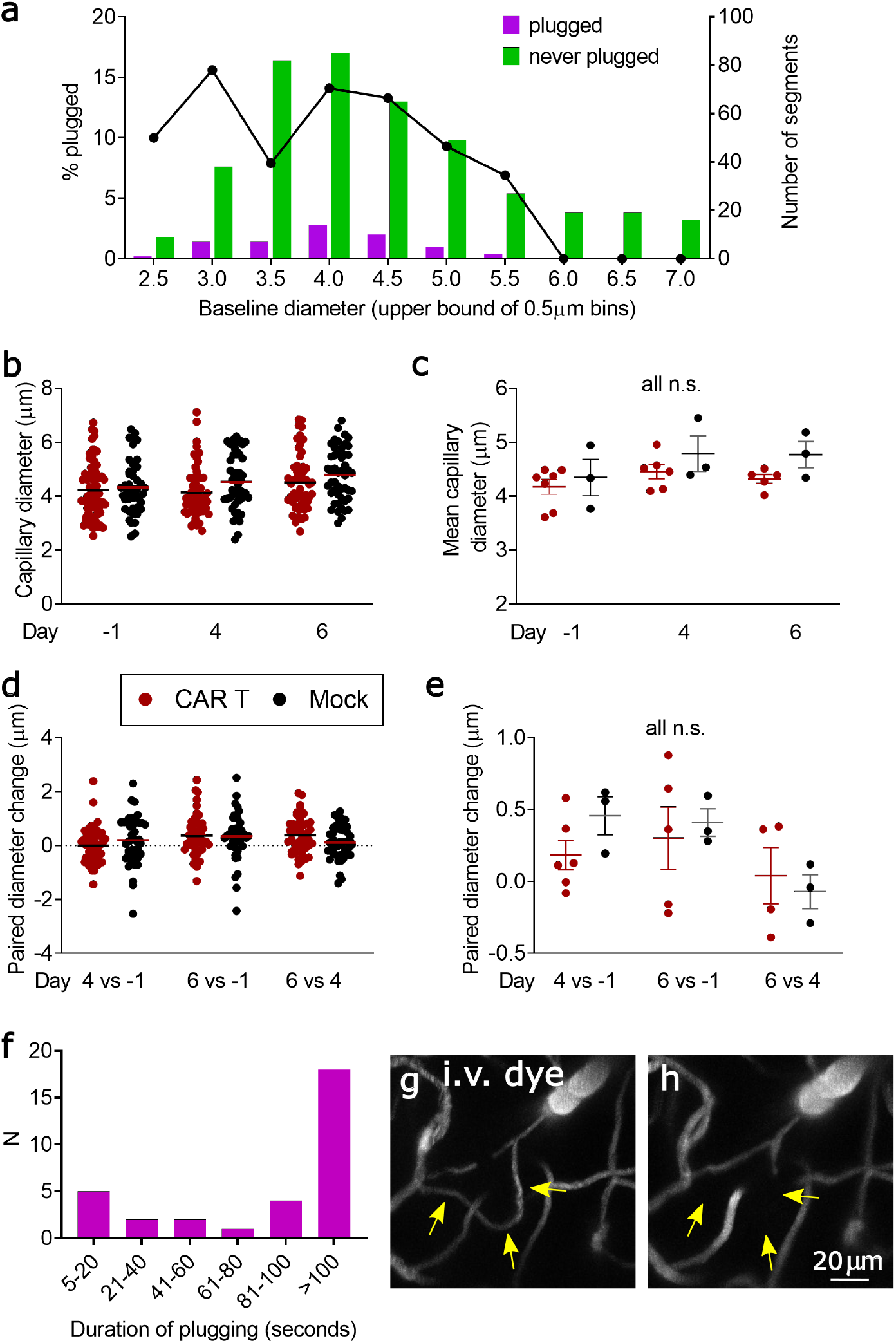
Diameter-dependence and kinetics of capillary plugging. **a**, baseline diameters (day-1) and likelihood of plugging on day 4 and/or 6. Left y-axis and line plot show the percentage of capillaries that had any plugging. The right y-axis and bar graphs show the raw counts of capillary segments in each size bin. Pooled data from 6 mice, each treated with 10 million CAR T cells. **b**, representative example of capillary diameters in one CAR T (red) and one mock (black) treated mouse. Each data point represents one capillary segment, all segments <7mm baseline diameter in 3 z-stacks were included. Line denotes the mean diameter. **c**, summary of mean capillary diameters from 3 separate experiments, each data point represents one mouse. Lines show mean and SEM, one-way ANOVA. **d**, representative example of diameter changes in individual capillary segments in the same animals as in **b**, each data point is one capillary segment, line denotes the mean. **e**, summary plot of diameter changes over time, each data point shows the mean from one mouse, lines show mean and SEM, one-way ANOVA. **f**, duration of plugging. Bars show the number of capillary plugs that persisted for the amount of time indicated on the x-axis. For plugs lasting <100s, both the beginning and the end of the stalling event were observed. Data pooled from 5 mice. **g, h**: rare loss of a capillary segment. Arrows show a flowing segment on day −1 (**g**), which no longer fills with dye on day 6 (**h**).

To better understand the kinetics of stalling, we obtained time series images of randomly selected capillary plugs. Of 32 cells observed for at least 100 seconds, 56% remained stationary the entire time (Fig. 6f). We reasoned that such prolonged interruption of blood flow might cause regression of the vessel, which would align with our findings of increased capillary remodeling in CAR T cell treated mice (Fig. 3). To test this, we identified 31 capillary segments that were plugged on day 4 and were well visualized again on day 6. Of the initially plugged segments, 74% were recanalized and flowing on day 6, 26% remained plugged, and none had regressed. When considering all capillary segments that we were able to follow from baseline imaging to day 6, whether later plugged or not, we were able to identify loss of any capillaries in only 2 of 7 CAR T cell treated mice (3/272 segments, 1.1% in one mouse; 1/243 segments, 0.4% in the other). We considered vessels as regressed if the entire segment had dropped out instead of having a cell-shaped filling defect (Fig. 6g,h). We found no loss of capillary segments in any of the control mice. These findings show that despite prolonged obstruction of capillaries, regression occurs only in a small fraction of segments during the acute phase of neurotoxicity.

## Discussion

We have developed a mouse model which recapitulates key aspects of clinical CAR T cell neurotoxicity – behavioral changes, association with CRS, and injury to cerebral microvessels. Using *in vivo* two-photon imaging, we discovered that plugging of cortical capillaries by leukocytes is a putative mechanism underlying neurotoxicity.

In our model, neurotoxicity begins during the first week after CAR T cell infusion, concurrently with CRS or very soon thereafter. Similarly, patients receiving CD19-directed CAR T cells typically develop neurotoxicity within days of the onset of CRS, often before CAR T cells are detectable in large numbers in the peripheral blood^8–10^. This may be explained by the fact that cytokine elevations are highest during the initial expansion of CAR T cells *in vivo*^8,16^, or possibly by a model where the presence of very few CAR T cells is sufficient to trigger a monophasic neurologic injury. This close temporal association of neurotoxicity differs from a prior model of CD19-CAR toxicity, where human hematopoietic stem cells and human CAR T cells were engrafted into immunodeficient mice transgenic for human hematopoietic growth factors (NSG-SGM3)^18^. Here, neurologic abnormalities did not develop until 4 weeks after CAR T cell infusion, when CRS had already resolved. In addition, histopathologic changes in this model consisted of meningeal infiltration of human macrophages, but no cerebrovascular changes were shown. These differences may be due to the fact that our syngeneic model allows murine immune mediators to interact with the murine neurovascular unit, setting into motion a signaling cascade that results in increased immune cell adhesion to the endothelium. Building on prior syngeneic models^26^, we were able to induce toxicity by increasing the doses of CAR T cells and lymphodepleting chemotherapy. It remains unknown why toxicity is dose dependent, since lower doses of CAR T cells can successfully engraft, expand, and eradicate tumors^20^. Higher doses may allow more rapid CAR T cell proliferation and higher cytokine release, and indeed we noted a correlation between cytokine levels and neurologic toxicity in our model.

We found that over 10% of cortical capillaries were obstructed by adherent leukocytes during acute neurotoxicity. This putative mechanism of neurologic injury has been previously described in a mouse model of Alzheimer’s disease, where obstruction of only 1.8% of capillaries by stalled neutrophils was correlated with neurologic dysfunction^36^. The authors used mathematical modeling to show that occlusion of 2% of capillaries was sufficient to reduce overall cerebral blood flow. While the depth of two-photon imaging only allows us to visualize the microvasculature of the cortex, we found that patchy hypoxia by hypoxyprobe labeling extended into deeper brain regions, including the hippocampus and thalamus. This suggests that capillary stalling is not restricted to the cortex. Regional differences in blood flow may account for the fact that some mice had high plugging but a relatively benign neurologic exam, and vice versa.

We propose that capillary plugging is the result of increased expression of adhesion molecules on the endothelium and/or leukocytes in response to proinflammatory cytokine signaling. If the plugging was a purely mechanical process caused by increased size or stiffness of circulating leukocytes, we would have expected an inverse relationship of capillary size and likelihood of plugging, which we did not find. As additional evidence for increased adhesiveness between leukocytes and endothelium, we noted high levels of leukocyte rolling and adhesion in larger venules that were too large to be obstructed. Targeting of adhesion molecules may provide a therapeutic modality for preventing or stopping neurotoxicity^38^. Additionally, if one specific cell type is responsible for the plugging, selective ablation may be helpful, while taken care to preserve the therapeutic potential of CAR T cells. For example, in mouse models of ischemic stroke, over 50% of capillaries in the core and penumbra of the stroke were plugged by neutrophils, erythrocytes, or platelet rich clots, and the plugging could be ameliorated by antibody-mediated ablation of neutrophils^39,40^. Further identification of the stalling cells in our model by *in vivo* two-photon microscopy will require strong, cell-type specific fluorescent labeling. Unfortunately, further distinction between CAR-expressing and CAR-negative T cells was not possible with the antibody against our Thy1.1 transduction marker as it did not yield sufficiently bright signal for *in vivo* imaging. Stable fluorescent labeling can also be accomplished by adoptive transfer of genetically modified cells, or the use of transgenic mice as T cell donors. The use of transgenic or knockout mice in syngeneic CAR T models is complex, since extensive backcrossing is required to achieve a uniform genetic background to avoid graft-versus-host disease^23,41^.

In addition to in-vivo capillary plugging, we found histologic evidence of injury to the neurovascular unit, including loss of coverage by capillary pericytes, increased capillary regression, and microhemorrhages. Understanding the relationship between these findings will require further studies. Based on the frequency of string capillaries observed on histology (249 per mm^3^), we would have expected to find approximately 2 regressed capillaries per z stack volume we had examined *in vivo*. In fact, we found much fewer examples of regression of capillary segments, which suggests that either *in vivo* imaging underestimated the number of regressed vessels, or that string capillaries represent a different biological entity. A role for direct on-target toxicity by CD19-CAR T cells was suggested by a recent study which noted expression of CD19 in fetal human brain vascular mural cells by single-cell RNAseq^42^. This mechanism is plausible, as destruction of pericytes and other vascular mural cells by CAR T cells could lead to compromise of the blood-brain-barrier or changes in blood flow^43,44^. However, although we found reduced coverage of capillaries by pericyte processes, the number of pericytes was unchanged in CAR T treated mice. Additionally, single-cell gene expression analysis of the mouse neurovascular unit shows low to undetectable CD19 expression in pericytes or vascular smooth muscle cells^45,46^. We did observe retraction of pericyte processes in mice with severe behavioral neurotoxicity, the mechanism and consequences of which remain to be explored. Other mechanisms of injury may include cytokine-mediated alterations of neurovascular unit integrity^47^, or direct insults to the endothelium by stalled immune cells. This may be similar to what occurs in during microvascular leukostasis caused by circulating leukemic blasts, which can be associated with microhemorrhages even in the absence of overt coagulopathy, indicating that the focal stalling causes loss of endothelial integrity^48^. Since all the capillary obstructions were CD45.2+ leukocytes and not platelet rich thrombi, our model does not support a role for thrombotic microangiopathy in the development of CAR T neurotoxicity^49^.

In summary, we have found evidence of neurovascular unit injury and leukocyte plugging of the cerebral microvasculature in a mouse model of CD19-CAR T cell therapy. This model can serve as a novel platform for dissecting the molecular mechanisms of CAR T cell toxicity, and may be useful in the development of safer cancer immunotherapies.

## Methods

### Animals

5-8 week old wild-type BALB/c mice were used for all experiments. Male and female mice were used in equal numbers. Mice were obtained from Jackson Labs (Bar Harbor, ME) and housed in specific pathogen free facilities. After CAR T cell treatment, ear temperature and weight were recorded daily, and mice received 10ml/kg normal saline subcutaneously daily. All experiments were approved by the Seattle Children’s Research Institute Animal Use and Care Committee.

### CAR T cell generation

The CAR construct (gift from Juno Therapeutics) consists of an anti-murine-CD19 single chain variable fragment (scFv) derived from the 1D3 hybridoma^21^ and murine intracellular 4-1BB and CD3ζ costimulatory domains, followed by a T2A sequence to coexpress the transduction marker Thy1.1. The construct was cloned into a MMLV backbone, and gammaretrovirus was produced by transient calcium phosphate transfection of a Phoenix-ECO producer line (ATCC). CAR T cells were generated per published protocol^50^. Briefly, T cells were collected by mechanically dissociating spleens and lymph nodes of wild type BALB/c mice, followed by MACS selection with CD90.2 beads (Miltenyi Biotec). T cells were then stimulated with CD3/CD28 Dynabeads (ThermoFisher) (days 1-4), murine IL-2 (Peprotech) (days 1-3) and murine IL-15 (Peprotech) (day 4) in RPMI supplemented with 10% fetal bovine serum, penicillin/streptomycin, β-mercaptoethanol, and sodium pyruvate. On days 2 and 3, cells were spinoculated with viral supernatant at 2000xg for 1 hour on retronectin (Takara) coated plates. Transduction efficiency was verified by flow cytometry for Thy1.1. Mock transduced T cells underwent the same transduction protocol, while replacing the viral supernatant with media during the spinoculation step.

### CAR T cell treatment

To reduce rejection of CAR T cells, mice received lymphodepleting chemotherapy with 250mg/kg cyclophosphamide via a single i.p. injection (Day −1). 24h later (Day 0), 5-10 x 10^6^ CAR T cells or mock transduced T cells per animal were infused via the lateral tail vein. Mice were monitored via daily weight and temperature measurements, and both controls and CAR T treated animals received daily subcutaneous normal saline or lactated Ringer’s solution boluses (20mL/kg). Blood for analysis was collected via chin bleed, retroorbital puncture under sedation, or post-euthanasia.

### Flow cytometry

Spleens were removed and mechanically dissociated through a 70μM filter. Blood was collected in EDTA tubes, RBCs lysed and lymphocytes collected via density gradient centrifugation in lymphocyte separation media. Mice were then perfused transcardially with ice-cold PBS to remove intravascular blood, and brains were dissected out. To recover brain infiltrating leukocytes, we used mechanical homogenization and density gradient centrifugation^51,52^. Brains were homogenized in RPMI in a 5mL Dounce homogenizer, and strained through a 70μm filter. Cells were suspended in 30% percoll in RPMI, and layered over a 70% percoll solution, followed by spinning at 500xg for 30 minutes at room temperature without brake. The myelin debris that collects at the top was removed, and the 30%/70% percoll interphase layer containining leukocytes was collected. Cells were then resuspended, labeled on ice with fluorescently conjugated antibodies against mouse CD45, CD3, CD19, Thy1.1, and viability dye (Biolegend), and analyzed on an LSR Fortessa or Novocyte flow cytometer.

### Cytokine analysis

Cytokine concentrations (CXCL1, IFNγ, IL-1β, IL-2, IL-4, IL-5, IL-6, IL-10, IL-12p70, and TNF) in serum and brain lysates were measured using the V-PLEX Proinflammatory Panel 1 Mouse Kit (Mesoscale Discovery) according to manufacturer’s instructions.

### Histologic analysis

Fluorescently labeled dextrans (5%, 50μL) (Thermo Fisher) were injected retroorbitally under isoflurane anesthesia 30 minutes prior to euthanasia. Mice were euthanized with CO_2_ and transcardially perfused with ice-cold PBS followed by 4% PFA. Brains were fixed in 4% PFA overnight, cryoprotected in 15% and 30% sucrose and cut into 20-50μm coronal sections. For hemorrhage counts, we analyzed H&E stained 30μm sections, which were taken every 1mm from the anterior brain excluding the olfactory bulbs, to the end of the cerebellum. The hemorrhages per volume analyzed were then normalized to a total brain volume of 500mm^3^ ^53^.

### Immunohistochemistry

20-30μm slide-mounted or 50μm floating PFA-fixed brain sections were incubated in blocking solution (10% donkey serum + 0.1% Triton X-100 in PBS) for 1 hour, then at 4C overnight with primary antibodies in 1% donkey serum + 0.1% Triton X-100 in PBS, washed in PBS, incubated with secondary antibodies for 2h at room temperature, counterstained with DAPI and mounted in Fluoromount G (Southern Biotech). Primary antibodies: Claudin-5 (mouse, clone 4C3C2, ThermoFisher cat# 35-2500, 1:100), Iba1 (goat polyclonal, Abcam cat# ab5076, 1:1000), laminin (rabbit polyclonal, Novus Biologicals cat# NB300-144, 1:50), CD13 (rat, clone 123H1, MBL cat# M101-3, 1:100). Slides were imaged on a Zeiss LSM 710 confocal microscope at 20x-63x magnification. Tissue from different treatment conditions was processed side by side and imaged at the same settings. The Fiji image processing package^54^ was used for analysis. All analysis was performed blinded to treatment condition. For IgG fluorescence quantification, 20μm sections were incubated with fluorescently conjugated antibodies directed either against mouse IgG, or the IgG of control species (rat, goat or rabbit) (all raised in donkey, Thermo Fisher) in PBS for 2h. The average brightness (arbitrary units) of immunofluorescence against mouse or control IgG was calculated for each imaging field. The median ratio of brightness of anti-mouse IgG staining to anti-control species IgG staining was calculated for at least 3 nonconsecutive brain sections per mouse. To normalize for differences in tissue treatment and incubation conditions between separate experiments, measurements were then expressed as % of mock treated control. For microglia counts and pericyte analysis, 50μm floating sections were used. Iba1 positive cells were counted manually on at least 3 separate z-stacks from different areas of cortex for each animal, and the median number of somata per mm^3^ was calculated. Pericytes were identified as protruding somata overlapping with DAPI nuclear staining and long extending processes along the vessel. We measured the length of laminin-labeled vessel segments, the length of pericyte coverage, the number of pericyte somata, as well as the number of string capillaries within a z-stack. We also subtracted background fluorescent to quantify the area of pericyte coverage. These values were then used to quantify pericyte coverage as a function of length and area, the number of pericyte soma per vessel length, and the number of pericyte soma and string capillaries per stack volume.

### Neurophenotype scoring

An observer blinded to treatment group conducted a daily 20-item neurophenotype exam. This was adapted from a screening exam for neurologic phenotypes in mutant mice^29^. Scored items include grooming, piloerection, respirations, extremity perfusion, eye opening, body posture, tail posture, spontaneous activity, visual orienting, walk on cage edge, whisker reflex, eye blink, ear reflex, startle reflex, balance on rod, climb onto rod, reach for target from suspended position, upper extremity grip strength, postural adjustment upon cage rotation, and unusual or stereotyped behaviors. Each item was scored 0=normal, 1=performed with limitations/mildly abnormal, or 2=unable to perform/severely abnormal, and the total daily score was determined by summing all items. Animals were habituated to the exam for 3 days in a row before any experimental manipulation was started.

### Open Field test

All mice were naïve to the test and were habituated to the behavior room for 30 minutes prior to testing. Testing was conducted under dim light, and the test box was thoroughly cleaned between animals. Mice were introduced into a 20×20×20cm box with a 10×10cm center area marked. Video was acquired for 15 minutes, and analyzed with Noldus software. Time spent in the center 10×10cm area, as well as total distance traveled, was quantified for the initial 5-minute interval after introduction into the test box.

### Echocardiograms

Serial echocardiograms were performed as previously described^55^. Briefly, during image acquisition mice were sedated with 1% isoflurane with 21% oxygen delivered via nose cone at 1L/min. ECG leads were placed for simultaneous ECG monitoring. Echocardiographic images were performed on a Vevo 2100 machine using a MS400 transducer (VisualSonics Inc, Toronto, Canada). M-mode measurements at the midpapillary level of the left ventricle were performed to measure endsystolic and enddiastolic diameters and calculate stroke volume.

### Cranial window preparation and anesthesia

We created a polished and reinforced thinned-skull window over the somatosensory cortex as previously described.^56,57^ Briefly, anesthesia was induced with 4% isoflurane and additional pain control provided with buprenorphine. Mice were maintained at 1-2% isoflurane for surgery. After removing the skin, the skull overlying the sensory cortex was thinned to translucency with a handheld drill. The area was covered with cyanoacrylate instant adhesive and a coverslip, and a custom head mount was attached using dental cement. Mice were allowed to recover for 24 hours prior to imaging. For imaging, anesthesia was induced with 4% isoflurane and mice were maintained at 1.5% while under the microscope. During *in vivo* imaging, all antibodies and intravascular tracers were delivered by injection into the retroorbital sinus. The blood plasma was labeled with 2mDa dextran conjugated to either FITC or Alexa 680 (5% in PBS, 20-50μl per injection). For *in vivo* leukocyte labeling, we injected either 50μl of freshly prepared Rhodamine 6G (0.1% in PBS), or monoclonal fluorescent conjugated antibodies (rat anti-mouse CD3, clone 17A2; mouse anti-mouse CD45.2, clone 104; all from Biolegend and used at 0.4mg/kg i.v.).

### *In vivo* two photon imaging

Imaging was performed on a Bruker Investigator microscope with Prairie-View imaging software, coupled to a Spectra-Physics InSight X3 tunable laser (680-1300nm) set to 800nm excitation. Green, red, and far red fluorescence emission was collected through 525/70nm, 595/50nm, and 660/40 bandpass filters, respectively. For the first imaging session at baseline, we collected an overview map of the entire window using a 4X (0.16 numerical aperture, NA) air objective (Olympus; UPLSAPO). The overview map was used to identify areas with unobstructed view of the cortical capillaries and to allow repeat imaging of the same areas at subsequent imaging sessions. We then switched to a 20X (1.0 NA) water-immersion objective (Olympus; XLUMPLFLN) and collected at least three 150μm deep image stacks per mouse. For antibody imaging, stacks were collected pre and post injection of antibody to control for any nonspecific fluorescence in capillary plugs.

### Two-photon image analysis

All analysis was performed in ImageJ/Fiji^58,59^. Every microvessel segment within an image stack was assigned a unique identifier using ImageJ’s multipoint function. Vessel labels were matched between baseline, day 4, and day 6 image stacks of the same areas. To count as nonflowing, a capillary had to have no movement of red blood cells on all image slices it was included, which were at least 5 slices for a typical capillary which took 6 seconds to acquire. We did not include brief capillary stalling in our counts, where flow can be slowed for <5 seconds by large white blood cells squeezing through a capillary. We then used the VasoMetrics ImageJ plugin^60^ to measure the diameter of each vessel segment at the full-width at half-maximum intensity. Segments of diameter over 7μm were excluded from analysis as they were likely to represent arterioles and venules.

### Immunostaining of pimonidazole adducts in hypoxic brain tissue

Fresh pimonidazole hydrochloride (Hypoxyprobe, Inc; HP3-100Kit) was prepared at 60μg/mL in sterile filtered PBS and injected at a dose of 60mg/kg into the retro-orbital vein under isoflurane anesthesia. Mice were removed from anesthesia and observed for 90 minutes to allow for pimonidazole adducts to form in any hypoxic tissue. We then euthanized the mice with COf_2_ and transcardially perfused with ice-cold saline followed by 4% paraformaldehyde. Brains were post fixed in 4% PFA for 24h, and cryoprotected in 15% and 30% (w/v in DI water) sucrose for 24h each. Using a cryostat (CM3050 S; Leica), we sectioned brains into 50μm thick coronal sections. Tissue sections were then incubated overnight at room temperature (RT) with anti-pimonidazole antibody (Pab2627 rabbit antisera; 1:100; Hypoxyprobe, Inc; HP3-100Kit) in a solution of 2% TritonX-100, 10% goat serum, and 0.1% sodium azide in PBS. The following day, sections were washed in a large volume of PBS (50–100 mL) for 30 minutes at room temperature before incubation with anti-rabbit Alexa Fluor 568 (A10042; 1:500; Invitrogen) and DAPI in PBS at RT for 2 hours. Finally, sections were washed one last time in a large volume of PBS for another 30 minutes. Sections were then mounted onto a glass slide and sealed with mounting medium (Fluoromount-G; SouthernBiotech) and cover glass.

## Acknowledgments

We would like to thank Eric Chadwick, Clay Patton, Rafael Ponce, Stephanie Busch, and Ruth Salmon for their work in the development of this mouse model.

